# Osmolarity-regulated swelling initiates egg activation in *Drosophila*

**DOI:** 10.1101/2021.02.16.431163

**Authors:** Anna H. York-Andersen, Benjamin W. Wood, Elise L. Wilby, Alexander S. Berry, Timothy T. Weil

## Abstract

Egg activation is a series of highly coordinated processes that prepare the mature oocyte for embryogenesis. Typically associated with fertilisation, egg activation results in many downstream outcomes, including the resumption of the meiotic cell cycle, translation of maternal mRNAs and cross-linking of the vitelline membrane. While some aspects of egg activation, such as initiation factors in mammals and environmental cues in sea animals, have been well-documented, the mechanics of egg activation in insects are less well understood. For many insects, egg activation can be triggered independently of fertilisation. In *Drosophila melanogaster*, egg activation occurs in the oviduct resulting in a single calcium wave propagating from the posterior pole of the oocyte.

Here we use physical manipulations, genetics and live imaging to demonstrate the requirement of a volume increase for calcium entry at egg activation in mature *Drosophila* oocytes. The addition of water, modified with sucrose to a specific osmolarity, is sufficient to trigger the calcium wave in the mature oocyte and the downstream events associated with egg activation. We show that the swelling process is regulated by the conserved osmoregulatory channels, aquaporins (AQPs) and DEGenerin/Epithelial Na^+^ (DEG/ENaC) channels. Furthermore, through pharmacological and genetic disruption, we reveal a concentration-dependent requirement of Trpm channels to transport calcium, most likely from the perivitelline space, across the plasma membrane into the mature oocyte.

Our data establishes osmotic pressure as the mechanism that initiates egg activation in *Drosophila* and is consistent with previous work from evolutionarily distant insects, including dragonflies and mosquitos, and shows remarkable similarities to the mechanism of egg activation in some plants.

## INTRODUCTION

Egg activation is a conserved process that prepares a mature oocyte for embryogenesis. It actuates many essential cellular processes including the resumption of meiosis, modification of the outer membrane, post-transcriptional regulation of maternal mRNAs and broad changes in the cytoskeletal environment [1–3]. This process requires a transient increase of intracellular calcium, often referred to as a calcium wave(s), with multiple waves observed in mammals and ascidians, compared to a single wave in *Xenopus laevis, Danio rerio* and *Drosophila melanogaster* [4–6].

Species variation is also documented in the initiation mechanism and source of calcium required for the cytoplasmic rise [1,4]. In vertebrates and some invertebrates, egg activation is dependent on fertilisation in which sperm entry introduces Phospholipase C enzymes generating a calcium efflux from the endoplasmic reticulum [2,7]. Comparatively, egg activation in other invertebrates can be independent of fertilisation and initiated by external factors [8]. For example, the ionic composition of the solution external to the oocyte is required in the starfish *Asterina pectinifera*, as chelation of sodium ions in seawater disrupted the resumption of meiosis [9,10]. While in the shrimp *Siconia ingentis*, egg activation requires the presence of magnesium ions in seawater [11]. Interestingly, in the stick insect *Catrausius morosus*, exposure of the oocyte to oxygen in the air results in the resumption of meiosis [12].

An alternative external cue of egg activation is the application of mechanical pressure on the oocyte plasma membrane exemplified by the eggs of the wasp *Pimpa turionellae*, which are activated when squeezed through a polythene capillary [13,14]. This physical stress is proposed to displace the maternal nucleus and result in the resumption of the cell cycle. Similarly, the eggs of *Drosophila mercatorum* are thought to be activated by the pressure from the genital ducts [12]. Tension in the plasma membrane can also be generated by a change in the osmolarity of the external solution (which we will refer to as ‘osmotic pressure’ henceforth). Prior to egg activation, the hypertonic environment in the ovaries is thought to maintain the oocytes in a meiotically-arrested state [15,16]. Subsequent entry of the oocyte into a hypotonic environment results in the egg activation of dragonfly, mayfly, turnip sawfly and yellow fever mosquito eggs [16–18]. For instance, upon entry into water, yellow fever mosquito oocytes undergo a visible darkening due to the increased production and cross-linking of the endochorion at egg activation [19,20]. Overall, physical pressure appears to be a conserved mechanism for initiating egg activation in many insects.

Similar to other insects, egg activation in *Drosophila melanogaster* is independent of fertilisation and occurs during the passage of the mature oocyte through the oviduct [21]. One model suggests that the pressure exerted by the oviduct on the oocyte upon entry initiates egg activation [8,22]. However, more recent work has shown that external pressure alone is not sufficient to trigger a calcium wave [5,23]. An alternative model proposes that osmotic pressure generated by uptake of oviduct fluid leads to the initiation of egg activation [15]. This is supported by observations that oocytes are visibly dehydrated whilst in the ovaries, but upon deposition appear turgid and hydrated [15,24]. Rehydration at egg activation can be recapitulated *ex vivo* through the addition of a hypotonic solution, known as Activation Buffer (AB), which when added to an isolated mature egg results in swelling and a single calcium wave [5,6,15]. This influx of calcium requires the Trpm mechanosensitive channel in the plasma membrane and results in the activation of Plc21C that sustains the wave [8,25,26]. Regulation of calcium entry was hypothesised to be related to distribution of the Trpm protein in the membrane, as calcium entry is often seen first at the poles. However, when observed using CRISPR-generated GFP-tagged Trpm, an even distribution of the protein across the plasma membrane was evident [27]. Therefore, the precise mechanisms of initiation and regulation of the calcium wave remain to be elucidated in *Drosophila*.

Here, we use live imaging in conjunction with novel physical manipulation, pharmacological disruption and genetics, to demonstrate the requirement of osmotically induced swelling for calcium entry and downstream events of *Drosophila* egg activation. We show that depletion of osmoregulatory machinery, including AQPs and DEG/ENaC channels, disrupts water homeostasis and egg activation. We provide further evidence that the movement of calcium ions into the egg is sensitive to levels of functional Trpm. Our data also argues that the external environment is not the source of calcium for the wave, but rather the ions are likely to originate from the perivitelline space. Together with other recent work in the field, our findings reveal that *Drosophila* egg activation has striking mechanistic similarities to other animals and even some plants.

## RESULTS

### Swelling is required for the initiation and propagation of the calcium wave

The likely initiation cues for the calcium wave at *Drosophila* egg activation include physical pressure applied on the posterior pole by the oviduct or the uptake of the fluid by the mature oocyte from the oviduct [8,15]. Our previous work has shown that physical pressure applied to the posterior pole is not sufficient to initiate the calcium wave [5]. This evidence, together with the observation that the mature oocytes are dehydrated whilst in the ovary but are turgid by the time they are deposited [15], suggests that swelling might play a role in the initiation and the propagation of the calcium wave at egg activation.

In addition to the initiation of the calcium wave, *ex vivo* dissected mature oocytes show an increase in oocyte volume, rounding of the oocyte poles and movement of the dorsal appendages following exposure to AB (Figure 1A). To test if this swelling is required for the initiation and propagation of the calcium wave, we blocked the ability of the egg to swell by placing the anterior pole in a plastic capillary with the posterior pole being exposed to oil (Figure 1B). Upon the addition of AB, the oil is displaced and the calcium wave initiated as normal. However, the wave did not propagate past the opening of the capillary (Figure 1B′). An uninhibited calcium wave would normally encompass the whole oocyte by 3.5 minutes [5]. However, in this case, the wave did not propagate until the egg was expelled from the capillary. When the whole oocyte was placed in the capillary, as expected, the calcium wave did not initiate upon the addition of AB (data not shown). This strongly suggests that swelling is required for the initiation and propagation of the calcium wave.

**Figure 1.**
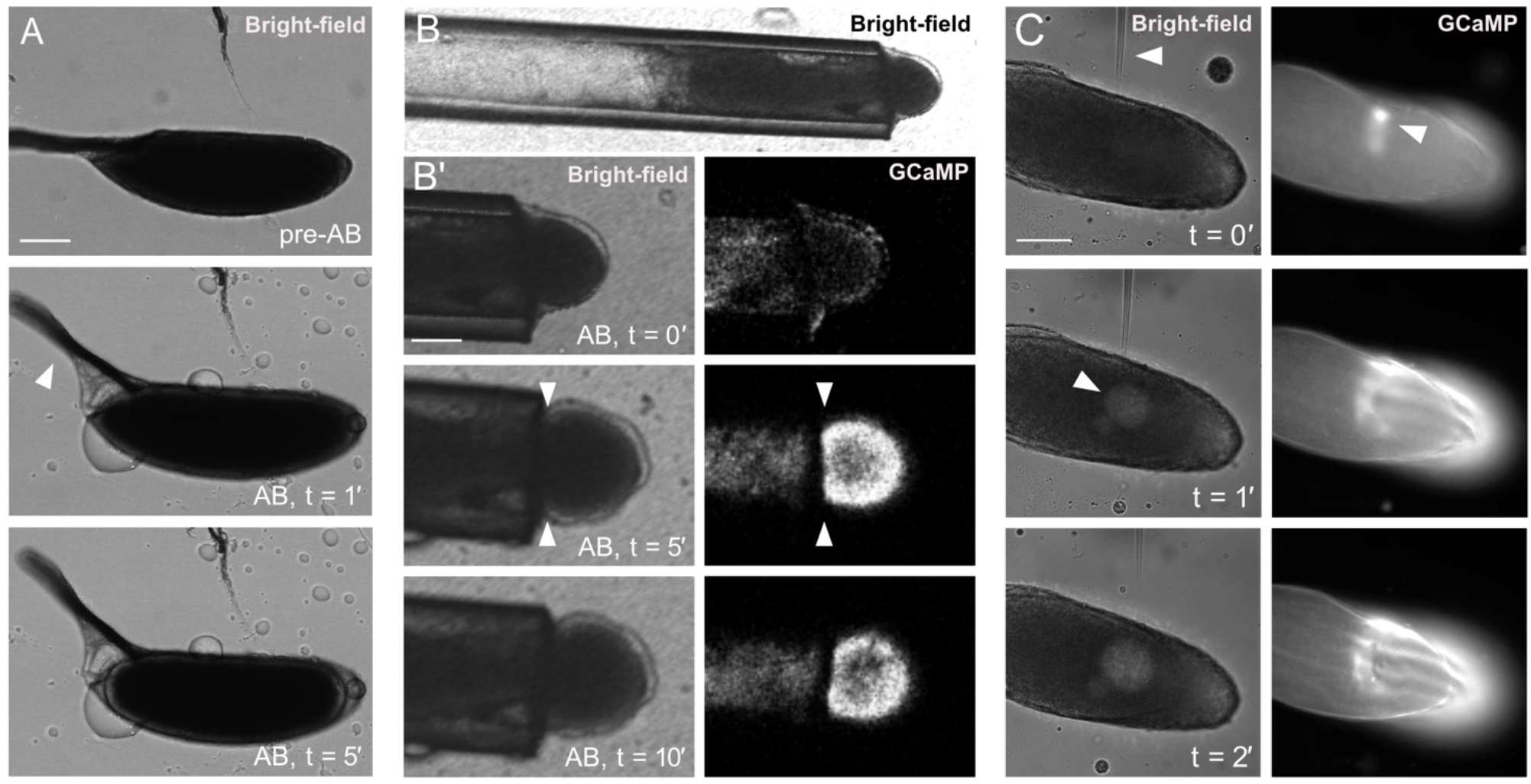
Egg chamber swelling is required for the initiation and propagation of the calcium wave. Time series showing *ex-vivo* mature egg chambers under bright-field (A,B,B′,C) and expressing *UAS-myrGCaMP5* (B,B′,C). Images represent a single plane. (A) Time series of a wild-type egg chamber pre- and post-addition of activation buffer (AB). Upon the addition of AB, the egg chamber undergoes swelling, the dorsal appendages rise (white arrowhead, t = 1′) and the poles become more rounded (white arrow, t = 5′). Circular droplets visible on the outside of the egg are oil that was not displaced by AB. Scale bar 100 μm. (B) Bright-field image of an egg chamber, expressing *UAS-myrGCaMP5*, placed in a 125 μm diameter tube. (B′) Time series of the same egg chamber in the tube, with the posterior pole exposed to AB. The calcium wave initiates normally but does not propagate past the tube opening (white arrowheads) (n = 15). Scale bar 60 μm. (C) Bright-field image of a mature egg chamber, expressing *UAS-myrGCaMP5*, injected with halocarbon oil. As the needle enters the oocyte, there is a calcium increase at the point of injection (white arrowhead, t = 0′). Injected oil is seen as a circle in the cytoplasm of the egg at t = 1′ (white arrowhead) and remains localised (t = 2′). Localised swelling results in an increase in calcium, which does not propagate (t = 2′) (n = 5). Scale bar 100 μm.

Our previous work has shown that local pressure or injection of calcium gives a localised calcium increase, but not a prolonged or broad calcium increase in a form of a wave [5,23]. To test if a localised internal increase in volume could induce a broad calcium event, we used a microneedle to inject halocarbon oil into a mature egg chamber mounted in halocarbon oil. Initial puncturing of the egg chamber resulted in a localised increase in calcium consistent with our previous results (Figure 1C, t = 0′). Injection of oil into the centre of the egg chamber resulted in a broad posterior calcium increase (Figure 1C, t = 1′, 2′). This response is noticeably different from previous experiments where oocytes were manipulated with a microneedle or had physical pressure applied. This data suggests swelling is necessary for the calcium propagation and is sufficient for the initiation of a broad calcium increase.

### Osmotic pressure initiates the calcium wave

In order to further test the function of swelling, we established a classification system that enabled us to categorise calcium events in the egg and quantify our data under different experimental conditions. We have classified the calcium increase exhibited as four distinct phenotypes: full wave, cortical increase, partial wave, or no wave. The most common phenotype is the full wave that initiates from the posterior pole and propagates across an entire oocyte. This is the standard event that we observe with *ex vivo* egg activation using AB. We do observe a small percentage of full wave phenotypes that initiate from the anterior pole. Different to a full wave, an increase in calcium can occur from multiple places around the cortex. This cortical increase phenotype was originally observed when egg chambers were exposed to distilled water [5]. These observations suggest that all parts of the egg have the capacity to allow calcium into the cell and that there is a regulatory mechanism to control calcium entry. In contrast, the partial wave phenotype describes the calcium wave that initiates from a pole but does not propagate across the entire oocyte and recovers prematurely. We do observe some egg chambers attempting to initiate waves multiple times, without successful propagation of calcium. Finally, the no wave phenotype describes an absence of a calcium increase anywhere in the egg for the length of the experiment. We observe this in a small percentage of eggs that are likely to have a major defect in development prior to dissection.

Our data indicates the requirement of swelling for the calcium wave to occur at egg activation (Figure 1B). One way the egg could undergo swelling is by exposure to a hypotonic solution, which would cause an influx of water and subsequently generate osmotic pressure within the mature oocyte. To test whether or not the uptake of water alone could act as an initiation cue for the calcium wave at egg activation, *ex vivo* egg chambers were treated with a sucrose and water solution (SW) of the same solute content as AB, measured in osmolarity (260 mOsm). The SW solution has no ions added and is very different from other buffers used to activate eggs. Sucrose is highly soluble in water and is neutrally charged making it suitable for varying the osmolarity of the solution. Upon the addition of SW, the egg chambers exhibited a similar proportion of the calcium wave phenotypes to AB (Figure 2A).

**Figure 2:**
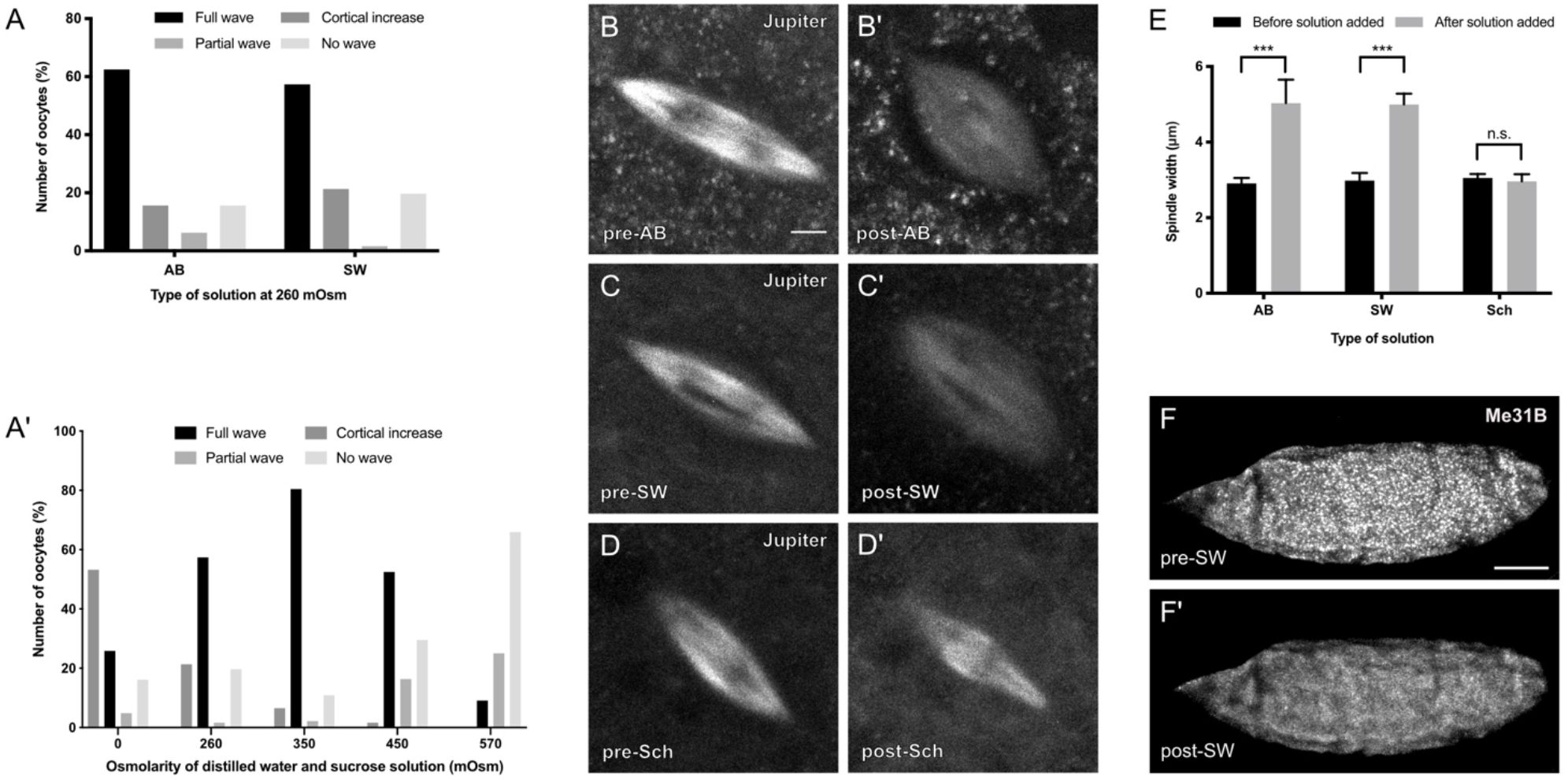
Osmotic pressure initiates the calcium wave and results in rearrangement of the meiotic spindle and P body dispersion. (A) Data showing activation buffer (AB) or sucrose and water (SW) of 260 mOsm results in a similar percentage of the calcium wave phenotypes when added to *ex vivo* egg chambers. (A′) The data shows the number of mature oocytes activated with SW only, with a range of osmolarities from 0-570 mOsm. The number of full waves increases from 0 mOsm, peaks at 350 mOsm and then decreases with higher osmolarities. The proportion of egg chambers that show a cortical increase peaks at 0 mOsm and then decreases with higher osmolarities. The proportion of partial waves increases with higher osmolarities. The proportion of no wave increases with higher osmolarities (n = 30 per osmolarity). This data was analysed statistically using Fisher’s exact test with P<0.05 considered significant. The proportion of full calcium waves observed at 350 mOsm is significantly higher (P<0.05) than full waves at all other osmolarities shown. The proportion of cortical increases observed at 0 mOsm is significantly higher (P<0.01) than cortical increases at all measured osmolarities shown. The proportion of partial waves observed at 570 mOsm is significantly higher (P<0.01) than partial waves at all measured osmolarities, except 450 mOsm. The proportion of no waves observed at 570 mOsm is significantly higher (P<0.001) than no waves at all other osmolarities shown. (B-D′) Mature egg chambers expressing *jupiter-mCherry* to visualise microtubules in the meiotic spindle. Before activation (pre) the spindle is in the shape of an ellipse, with dark regions in the middle where the DNA resides (B-D). Post-incubation images were taken 10 minutes after the addition of the solution. The spindle shows an increase in width following the addition of AB and SW (260 mOsm) (B-C′), however, the width does not change upon the addition of Schneider’s *Drosophila* Medium (Sch) (n = 15). Scale bar 2μm. Maximum projection 3 μm. (E) Graph showing a change in spindle width upon addition of AB, SW and Sch. The data was analysed statistically using an unpaired T-test with P<0.05 considered significant. The spindle shows a significant increase in width by 2.1 μm (a 1.7x increase) (P<0.001) upon the addition of AB or SW (260 mOsm). There is no significant change in width upon the addition of Sch (n = 15 per solution). (F-F′) Time-series of *ex vivo* egg chamber expressing *me31B::GFP* following the addition of SW (260 mOsm). P bodies appear as granular puncta pre-SW and disperse following the addition of SW, consistent with the addition of AB (n = 15). Post-incubation images were taken at 10 minutes after incubation of solution. Scale bar 60 μm. Maximum projection 40 μm.

To further test whether the osmolarity of an external solution is important for the initiation of an internal calcium increase, egg chambers were exposed to a single SW solution from a range of osmolarities. The highest percentage of full calcium waves was observed at 350 mOsm, with this percentage declining rapidly by 570 mOsm (Figure 2A′). The highest proportion of cortical increases was detected at 0 mOsm (Figure 2A′), consistent with predictions that an excessive volume increase cannot be regulated by the egg and results in an uncontrolled calcium increase. The partial and no wave phenotypes became more predominant with an increase in the osmolarity. This suggests that high osmolarity solutions do not increase the internal volume that is required for the egg to complete a calcium event. Together, these findings suggest that a controlled amount of water entering the egg is important for regulating a calcium event at egg activation.

### Osmotic pressure results in the resumption of the meiotic cell cycle

Previous work has shown that the addition of AB to mature egg chambers can initiate major cellular events associated with *Drosophila* egg activation, including the resumption of the cell cycle and P body dispersion [5,28]. In a non-activated oocyte, the meiotic spindle is parallel to the cortex and is observed near the base of the dorsal appendages at the anterior pole [28–30]. Upon egg activation, the spindle undergoes a morphological change within 10 minutes, marking the resumption of the cell cycle [28].

To address whether osmotic pressure alone results in this change, we used Jupiter-mCherry to label the meiotic spindle and exposed these mature egg chambers to SW solution (260 mOsm). Before exposure, the spindle is a narrow ellipse with dark regions in the middle where the DNA resides (Figure 2B,C,D). Upon addition of SW and AB (positive control) the spindle shows a significant increase in width (70%), which is indicative of spindle contraction at Anaphase I (Figure 2B′,C′,E). When treated with Schneider’s *Drosophila* Medium (negative control) the spindle did not undergo any detectable morphological change (Figure 2D′,E). Together, this supports the conclusion that water uptake and subsequent internal pressure is sufficient to initiate the Metaphase I-to-Anaphase I transition of the meiotic spindle.

To further verify the role of osmotic pressure we investigated the dispersion of P bodies, an established hallmark of egg activation [5,31]. When mature egg chambers expressing a conserved P body marker are exposed to SW (260 mOsm), we observe a normal dispersion phenotype (Figure 2F-F′). Taken together, this data suggests that an increase in internal volume caused by osmotic pressure triggers downstream events of egg activation.

### Water homeostasis is required for egg activation

The increase in internal volume observed in osmolarity experiments is regulated by water influx and efflux. To test if water homeostasis is required to regulate swelling in mature oocytes, we explored the role of the water-pore channels, AQPs, which are known to coordinate the movement of water molecules [32,33]. By adding copper sulfate (a broad AQP channel antagonist) into AB we do not observe a calcium wave (Figure 3A).

**Figure 3.**
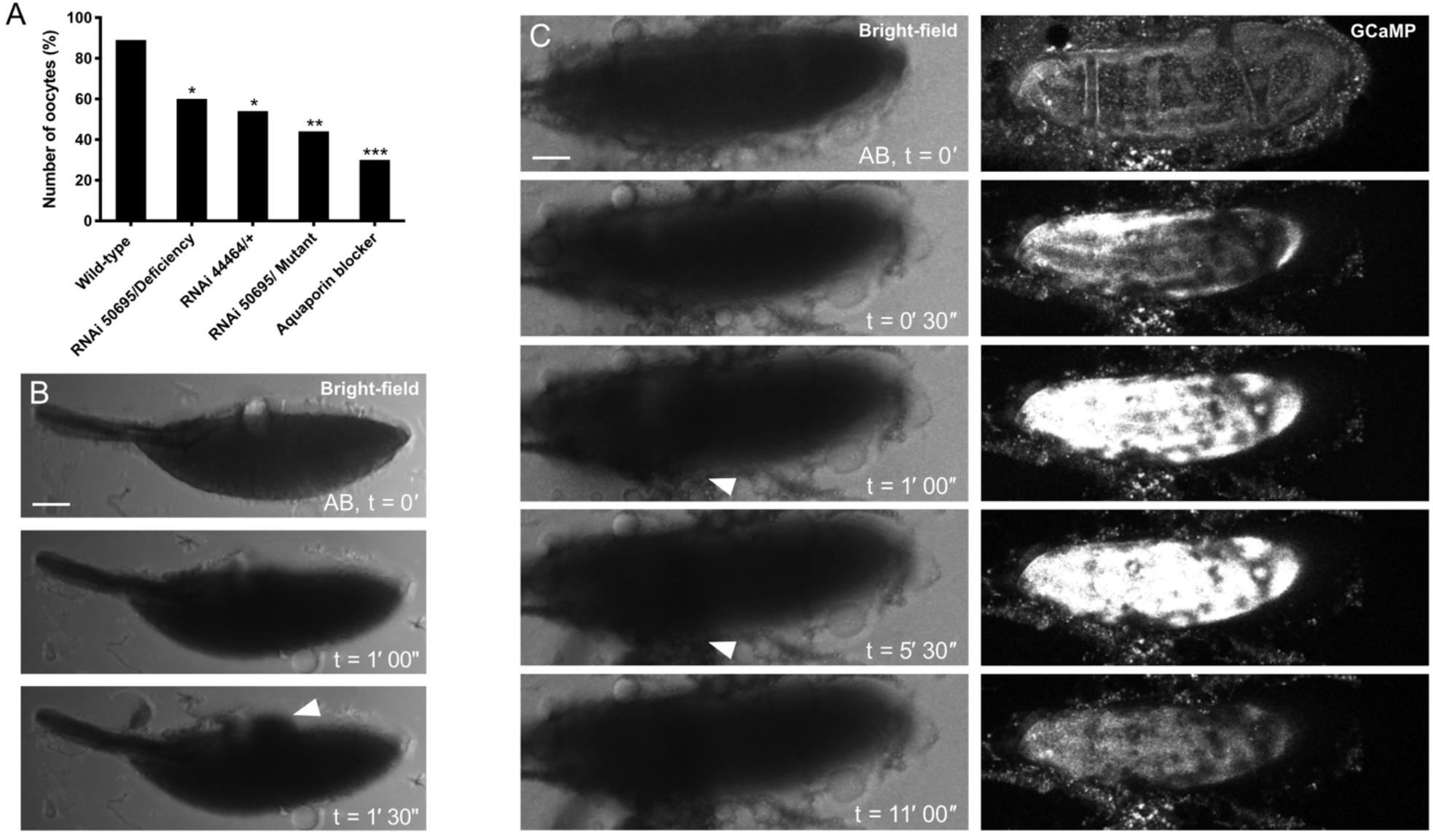
Water homeostasis is required for egg activation. (A) The data shows the presence of the calcium wave in Aquaporin depleted backgrounds. Aquaporin depletion was achieved through knockdown using BL50695 (germline) and BL44464 (germline and somatic) RNAi, deficiency (Df(2R)BSC160/Cyo), *prip* mutant (y1; P{SUPor-P}PripKG08662) and the broad Aquaporin channel antagonist copper sulfate. Upon the addition of AB, the number of oocytes with the calcium wave significantly decreased to 50% in the germline knockdown over the deficiency (n = 25, P<0.05) or mutant (n = 18, P<0.001). A similar significant decrease was also observed with only one copy knock-down of both somatic and germline Prip (BL44464) (n = 13, p<0.05). Addition of copper sulfate results in a significant decrease of waves to approximately 30% (n = 44, P<0.001). (B) Bright-field time series of mature egg chamber in an Aquaporin depleted background (RNAi 50695/deficiency). Upon addition of AB, 50% of the oocytes burst compared to 3% in the wild-type, with the cytoplasm leaking within 1 minute and 30 seconds (white arrowhead) (n = 120). Scale bar 60 μm. Maximum projection 40 μm. (C) Time-series of *ex vivo* mature egg chamber expressing *UAS-myrGCaMP5* and two copies of *ripped-pocket* RNAi following the addition of AB. The cortical increase appears within 30 seconds of the addition of AB, which is followed by the oocyte burst and cytoplasm leaking out in 74% of egg chambers (white arrowhead). The dark spots represent excess tissue and oil droplets. Scale bar 60 μm. Maximum projection 40 μm.

There is only one AQP channel, Prip, that is known to be expressed in the *Drosophila* ovarian tissue (*Drosophila* Fly Atlas). To investigate the role of Prip at egg activation we used knock-down tools in heterozygous deficiency or mutant backgrounds since the homozygous mutant was lethal. Upon the addition of AB, the number of oocytes showing a calcium wave significantly decreased in egg chambers expressing various AQP deficient backgrounds (Figure 3A). We further investigated the effect of Prip disruption by observing the time series of *ex vivo* activated eggs and found that half of these eggs rupture and leak cytoplasm shortly after the addition of AB (Figure 3B). Interestingly, some eggs were still able to initiate and propagate a calcium wave despite rupturing. This data is strongly suggestive of a subsequent requirement for Prip in mediating water homeostasis at egg activation.

Similar effects were observed in egg chambers expressing reduced levels of *ripped-pocket (rpk)*, a member of the mechanosensitive channels family DEG/ENaC known to be involved in transducing changes in osmotic pressure [34–36]. When activated, these egg chambers show a cortical calcium increase, rupture of the plasma membrane and leaking of the cytoplasm (Figure 3C). This phenotype is similar to when eggs are exposed to low osmolarity solutions (Figure 2A′), suggesting that *rpk* is required to mediate water entry. Together, this data suggests the role of AQP and DEG/ENaC channels is to coordinate optimal swelling and water homeostasis at egg activation.

### External calcium is not required for initiation and propagation of the calcium wave

In many animals, external and/or internal calcium is required for the calcium rise at egg activation [4]. To investigate the source of calcium at *Drosophila* egg activation, *ex vivo* mature egg chambers were treated with AB containing the calcium chelator BAPTA. These eggs exhibited typical swelling and a full calcium wave (Figure 4A), suggesting that external calcium from the surrounding solution is not required. To further validate this experiment we depleted internal calcium by pre-incubating egg chambers with membrane-permeable BAPTA-AM (with solubilising agent PF-127). The addition of AB with this chelator significantly reduced the number of calcium events (Figure 4A). Together, these findings point towards the source of calcium residing within the mature egg chamber.

**Figure 4.**
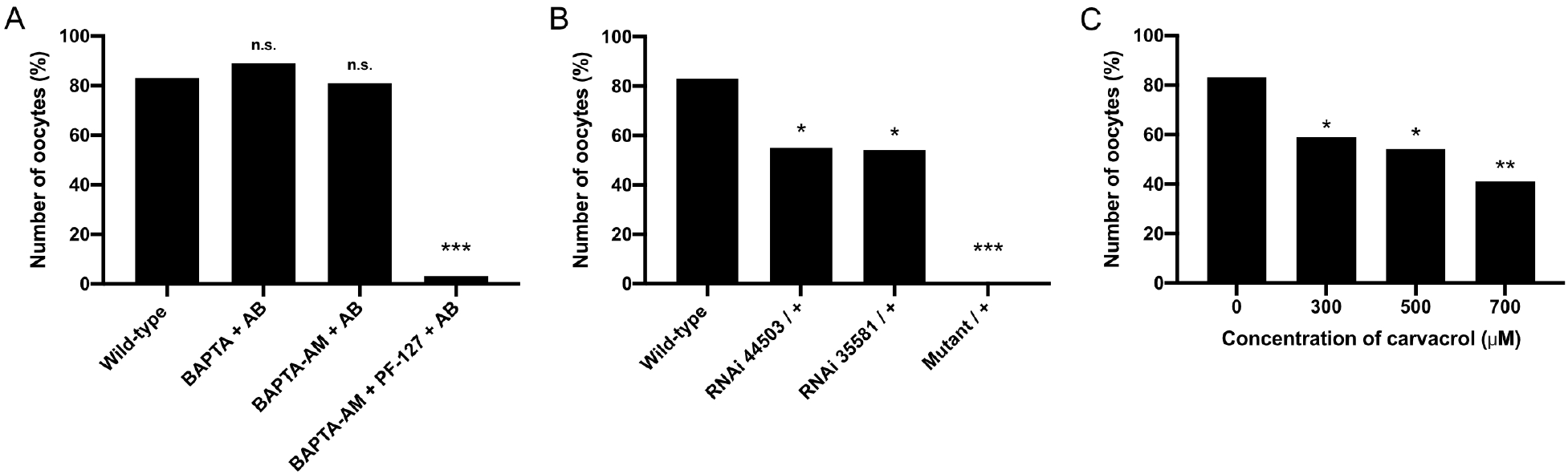
Internal calcium and Trpm channel are required for the calcium wave. (A) The graph shows the presence of the calcium wave in external or internal calcium-depleted backgrounds. Depletion of external calcium was achieved through the addition of calcium chelator BAPTA or BAPTA-AM in AB. Depletion of internal calcium was achieved through the addition of BAPTA-AM and PF-127 in AB. Upon the addition of BAPTA or BAPTA-AM in AB, there was no significant difference in the number of oocytes with the calcium wave (n = 19 and n = 57). The addition of BAPTA-AM and PF-127 in AB resulted in a significant decrease in the number of calcium waves (n = 34, P<0.001). Data was statistically analysed using Fisher’s exact test. (B-C) The graphs show the presence of the calcium waves in Trpm depleted backgrounds. Trpm depletion was achieved through knockdown using BL44503 (somatic and germline) and BL35581 (germline) RNAi, *trpm* mutant (y1 w67c23; P{EPgy2}TrpmEY01618/CyO) (B) and (C) the broad Trpm blocker carvacrol. Upon the addition of AB (B), the number of the oocytes with the calcium wave significantly decreased with only one copy knockdown of both somatic and germline Trpm (BL44503) (n = 96, P<0.01) and germline only (BL35581) (n = 35, P<0.05). A significant decrease in the number of the calcium waves was also observed in Trpm mutant background (n = 14, P<0.001). (C) A significant decrease in the number of the calcium waves was also observed with the addition of AB with the broad Trpm blocker carvacrol in a concentration-dependent manner of 300μM (n = 27, P<0.05), 500μM (n = 24, P<0.05) and 700μM (n = 24, P<0.01). Data was statistically analysed using Fisher’s exact test.

There are several potential internal calcium sources in the mature egg chamber, including the perivitelline space surrounding the mature oocyte [22]. The perivitelline space has been shown to consist of different ions, including calcium in the early *Drosophila* embryo [37]. However, it remains technically not possible to extract this fluid from the mature egg chamber due to the dehydrated morphology. One candidate, previously shown to be involved in coordinating the entry of calcium from the perivitelline space into the oocyte, is the mechanosensitive channel Transient Receptor Potential M (Trpm) [26].

To further investigate the role of Trpm, we utilised a transgenic line from the Berkeley *Drosophila* Genome Project which is a transposon P-element insertion in the 39th splice site which results in an imprecise deletion of three exons of Trpm [38,39]. Upon the addition of AB, these mature egg chambers swelled as expected, but fail to initiate a calcium increase (Figure 4B). We further tested the requirement of Trpm using a germline RNAi, which in heterozygous egg chambers resulted in a significant reduction in calcium waves (Figure 4B).

Together, this suggests that there could be a concentration-dependent response of Trpm. To test this hypothesis, egg chambers wild-type for Trpm were incubated with different concentrations of Carvacrol, a broad Trpm inhibitor [40]. The addition of AB with Carvacrol resulted in a significant decrease of the number of eggs with a calcium wave at a range of concentrations (Figure 4C). These data support the findings that Trpm is involved in regulating the entry of calcium into the mature oocyte at egg activation. Since Trpm channels are located in the cell membrane [26], it is likely that their role is to allow calcium from the perivitelline space to enter the oocyte at activation.

## DISCUSSION

### Model of *Drosophila* egg activation

In summary, our data shows that the calcium wave and characteristic downstream events associated with egg activation are initiated by osmotic pressure generated by the uptake of external fluid. We show that AQP and DEG/ENaC channels are required for mediating water homeostasis to withstand the rise in osmotic pressure during egg activation. We present complementary evidence that the Trpm channel is required for the influx of calcium, which we show is not supplied from a source outside of the egg chamber.

Together with previous work, our data supports the following model of *Drosophila* egg activation (Figure 5): (1) at ovulation, the meiotically-arrested mature oocyte passes into the lateral and then common oviduct; (2) the mature oocyte then takes up fluid due to the difference in osmolarity between the oviduct fluid and the ooplasm; (3) the increase in volume results in tension at the plasma membrane and dispersion of the cortical actin; (4) decreased density of cortical actin at the poles, prior to dispersion, primes these regions for calcium entry; (5) calcium enters the egg from the perivitelline space through the mechanosensitive Trpm channels in the plasma membrane; (6) starting at the posterior pole, further increase in intracellular calcium is relayed across the oocyte by the opening of the neighbouring Trpm channels via the dispersion of the cortical actin cytoskeleton at the lateral sides, resulting in the calcium wave propagation across the oocyte. (7) The calcium wave is then followed by an F-actin wavefront, which ensures the reorganisation of the actin cytoskeleton; (8) intracellular calcium returns to basal levels, likely through channels that transport calcium back into the perivitelline space. Collectively, the single calcium wave prepares the oocyte for pronuclear fusion and embryogenesis.

**Figure 5.**
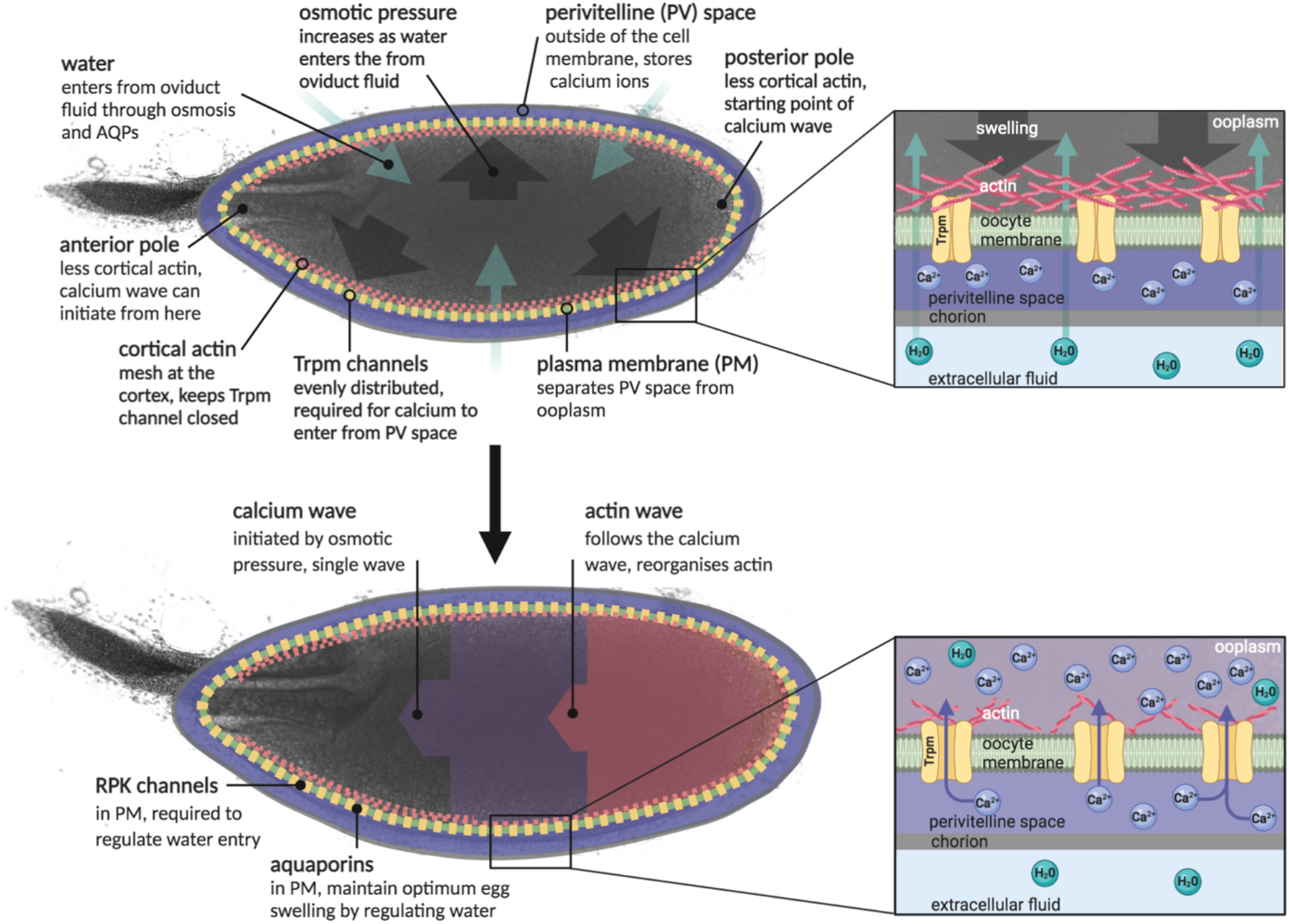
Model of *Drosophila* egg activation. Essential components and processes of *Drosophila* egg activation are outlined in the panels. A comprehensive description of the model is included in the discussion. Created with BioRender.com.

Osmotic pressure is a common mechanism for a volume increase and a rise in intracellular calcium levels, exemplified by intestinal epithelial cells, human osteoblast-like cells, rat astrocytes and cancer cell lines [41–44]. It is hypothesised that cells sense an increase in cell volume via intracellular solute, membrane-bound and/or cytoskeletal sensors [45]. The application of osmotic pressure seems to be a conserved initiation cue for egg activation in insects. Previous work has shown that the immersion of the oviposited mature oocytes of the yellow fever mosquito into water can resume oocyte development [46]. Similarly, for oocytes of the turnip sawfly and the malaria vector mosquito, egg activation can be initiated by placing the oocytes into water [16,17].

*Drosophila* is currently the only example of an insect in which the mature oocytes have been shown to exhibit an increase in intracellular calcium in response to the addition of hypotonic solution [5,6]. Our results presented here show that osmotic pressure acts as the initiation cue of the calcium wave at *Drosophila* egg activation.

### AQP and Rpk requirement in water homeostasis

It is essential to regulate cellular volume in response to changes in osmotic pressure. This is often achieved by AQPs, a conserved channel known to control the influx and efflux of water during cellular processes, including cell migration, neuroexcitation and epithelial fluid transport [32]. In *Drosophila*, our findings show the AQP Prip is required to maintain an optimal volume change at egg activation. In a *Prip* depleted background we observed eggs initially swelling but rupturing shortly after. This phenotype suggests that Prip is required to remove water from the oocyte as the egg swells during egg activation.

In addition, we show that the depletion of the DEG/ENaC channel, *Rpk*, also results in oocytes rupturing when activated. We propose that Rpk mediates optimal swelling through interactions with the cortical actin cytoskeleton, which we have previously shown to be re-organised at egg activation [23]. This hypothesis is supported by (1) co-immunoprecipitation studies in MDCK cells in which DEG/ENaC channels bind F-actin via the COOH terminus of α-ENaC and (2) mechanical pressure experiments that activate DEG/ENaC channels resulting in the stiffening of the cortical actin in vascular endothelial cells [47,48]. We therefore propose that Rpk is stabilising the cortical actin to withstand the increase in volume at activation.

### Role of osmotic pressure and TRP channels at egg activation

Recent work on germ-line knockout mutants in *Drosophila* have established the requirement of mechanosensitive Trpm channels in mediating the calcium influx at egg activation [26]. We corroborate this requirement using different mutants, RNAi and pharmacological disruption. Our data supports a model in which osmotic pressure generates tension in the plasma membrane and the cortical actin resulting in the opening of Trpm channels and subsequent calcium entry. Interestingly, the mammalian homolog TRPM3 is also activated in HEK293 cells by the application of a hypotonic solution, resulting in an intracellular calcium increase [49]. Similarly, in mammalian sensory neurones, TRPV4 and TRPV1 respond to changes in osmotic pressure [50–53].

Calcium entry mediated by TRP channels appears to be a conserved mechanism in the eggs of many animals. This was first shown in *Xenopus* oocytes where a mechanical stimulus resulted in the opening of TRPC1 [54]. More recently, mouse oocytes have been shown to require TRPV3 for the calcium intracellular increase and were affected by overexpression and the application of 2-APB [55]. In addition, TRPM7 was also shown to be essential for the calcium influx at mouse egg activation [56]. Finally, in *Caenorhabditis elegans* loss of the TRP3 channel resulted in a failure to show a calcium rise at egg activation [57]. Taken together these examples highlight a conserved role of TRP channels in mediating successful egg activation through calcium entry.

### The source of calcium at *Drosophila* egg activation

Calcium waves at egg activation can be mediated by intracellular and/or external calcium sources [4]. In *Drosophila*, Trpm regulates calcium entry across the plasma membrane suggesting that the calcium source is external to the oocyte. Paradoxically, we also show that external calcium is not required for a wild-type calcium wave. We argue that this data is compatible and point to the perivitelline space, situated between the oocyte plasma membrane and the vitelline membrane, as the calcium store. The composition of the perivitelline space in the egg chamber is currently unknown. However, in the early embryo, it has been shown to consist of many ions including calcium [37]. Our work supports a model where the perivitelline space is pre-loaded with calcium during oogenesis which enters through Trpm channels when the egg swells. This model is supported by our data showing that the injection of oil (devoid of calcium) is sufficient to induce a calcium rise in the oocyte.

### The *Drosophila* calcium wave is an example of a “slow” calcium wave

While calcium waves can be classified by the source of ions, they can alternatively be compared based on how fast they propagate [58]. In most animals, calcium waves at egg activation are classified as fast, travelling at ∼10-30 μm/sec [59]. However, some wave(s) propagate at ∼0.2-2 μm/sec and are classified as slow. This includes calcium influx at egg activation in maize eggs which propagates at 1.13 μm/sec and interestingly, requires mechanosensitive channels [60–62]. This is very similar to observations in *Drosophila*, where mechanosensitive channels and the actin cytoskeleton are required for a slow wave that propagates at ∼1.5 μm/sec. In fact, the general mechanism for a slow calcium wave [56] is strikingly similar to what we propose is occurring at *Drosophila* egg activation. Overall, aspects of the calcium wave, and more broadly egg activation, in *Drosophila* appear to be conserved with a variety of other organisms. Further analysis in flies will likely show even more similarities and inform our overall understanding of egg activation in all species.

## MATERIALS AND METHODS

### Fly stocks

The following fly stocks were used: *UASt-myristoylated(myr)-GCaMP5*); *matα-GAL4::VP16* (BL7063) and *UASp-GCaMP3* [6]; *tub-GAL4VP16* (Siegfried Roth); *jupiter-mCherry* (Paul Conduit); *me31B::GFP* [63]; *ripped-pocket RNAi* (P{TRiP.HMS01973}attP40, BL39053); *trpm mutant* (P{EPgy2}TrpmEY01618/CyO, BL15365); *trpm RNAi* (BL35581 and BL44503); *prip mutant* (P{SUPor-P}PripKG08662, BL14750); *prip RNAi* (P{TRiP.GLC01619}attP2, BL44464); *prip RNAi* (P{TRiP.HMC03097}attP40, BL50695); deficiency (for *prip*) (*Df(2R)BSC160/CyO*, BL9595). Stocks were raised on standard cornmeal-agar medium at 21°C or 25°C. For dissection of mature oocytes, mated females were fattened on yeast for 48 hours at 25°C.

### Reagents

BAPTA (Sigma-Aldrich) was used at a final concentration of 10 µM; BAPTA-AM + PF-127 (Sigma-Aldrich) was used at a final concentration of 30µM; Carvacrol (Sigma-Aldrich) used at 300-700µm. For the above reagents, standard preparation protocols were used as according to Sigma-Aldrich.

Activation Buffer (AB) containing 3.3 mM NaH_2_PO_4_, 16.6 mM KH_2_PO_4_, 10 mM NaCl, 50 mM KCl, 5% polyethylene glycol 8000, 2 mM CaCl_2_, brought to pH 6.4 with a 1:5 ratio of NaOH:KOH [15]; Gibco Schneider’s *Drosophila* Medium (Thermo Fisher); Series95 halocarbon oil (KMZ Chemicals); EZ-Squeeze tube 125 µM (Cooper Surgical). For osmolarity experiments, sucrose (Sigma-Aldrich) was directly dissolved into distilled water and the osmolarity was measured using an osmometer (Löser).

### Preparation of mature oocyte for live imaging

Mature oocytes were dissected from the ovaries from fattened flies using a probe and fine forceps [64]. Dissected oocytes were placed in series 95 halocarbon oil (KMZ Chemicals) on 22 × 40 coverslips, aligned parallel to each other to maximise the acquisition area for imaging, left to settle for 10 minutes, and incubated in solution *ex vivo* [64].

### Imaging

Time-series were acquired with an inverted Leica SP5, under 20x 0.7NA immersion objective. The Z-stacks were acquired at 2 μm steps from the first visible plane to 40 μm deep. The Z-stacks were presented as maximum projections of the 40 μm unless stated otherwise.

### Oil injection

Preparation for microinjection was carried out with a Femtotips II microinjection needle (Eppendorf) and a gas pressure injection system were used to inject oil into the Stage 14 egg chambers [31]. Imaging was performed simultaneously with injection on a DeltaVision wide-field microscope (Applied Precision) using a 20x 0.75NA numerical aperture.

### Quantifications and analysis

The calcium wave data was analysed statistically using Fisher’s exact test with P-values (P<0.05 considered significantly different) [6]. The spindle dimensions were quantified and statistically analysed using an unpaired T-test with P<0.05 values showing significant difference. The number of asterisks represents the P-value: (*) P≤ 0.05; (**) P≤ 0.01; (***) P≤ 0.001.

## ACKNOWLEDGEMENTS

We are grateful to Richard Parton for experimental and technical advice; Mariana Wolfner, Matthias Landgraf, Howard Baylis, José Casal, Peter Lawrence for feedback, discussions and advice; Richard York-Weaving for feedback on the manuscript; the Zoology Imaging Facility and Matt Wayland for assistance with microscopy; Siegfried Roth, Paul Conduit and Mariana Wolfner for fly stocks; and funding from the University of Cambridge ISSF grant number 097814 (to TTW), Department of Zoology Balfour Studentship (to AHYA), BBSRC DTP studentship (to BWW), BBSRC DTP studentship (ELW), and Sir Isaac Newton Trust Research Grant (Ref 18.07ii(c)).

